# Application of Different Types of Lactic Acid Bacteria Inoculant on Ensiled Rice Straw; Effects on Silage Quality, Rumen Fermentation, Methane Production and Microbial Population

**DOI:** 10.1101/612556

**Authors:** Ehsan Oskoueian, Saeid Jafari, Reza Noura, Mohammad Faseleh Jahromi, Goh Yong Meng, Mahdi Ebrahimi

## Abstract

Bacterial inoculants are known to improve quality of silage. The objectives of the present study were to evaluate the effects of different types of lactic acid bacteria (LAB; *L. plantarum*, *L. salivarius, L. reuteri, L. brevis* and *S. bovis*) inoculation (10^6^ g^−1^ DM) on rice straw silage quality and to examine these effects on ruminal fermentation characteristics, digestibility and microbial populations in an *in vitro* condition. Inoculated rice straw was ensiled for 15 and 30 days. For *in vitro* study, rumen liquor was obtained from two rumen fistulated mature cows fed on mixed forage and concentrate at 60:40 ratio twice daily. Inoculation of LAB improved (P˂0.05) the rice straw silage quality such as increased dry matter and crude protein contents, decreased pH and butyric acid, and increased propionic acid and LAB contents especially after 30 days of ensiling. Results from *in vitro* study revealed that addition of LAB to the rice straw silage improved fermentation characteristics such as increased total volatile fatty acids and dry matter digestibility (P˂0.05). LAB treatments also decreased methane production and methane/total gas ratio after 15 and 30 days of ensiling. From the rumen microbial population perspective, cellulolytic, and fungal zoospores were enhanced while protozoa and methanogens were decreased by the LAB treatments. Based on these results, it could be concluded that inoculating rice straw silage with LAB (especially for *L. plantarum* and *S. bovis*) improved silage quality, rumen fermentation parameters and microbial populations *in vitro*. However, *in vivo* studies need to confirm those effects.

## Introduction

Use of agricultural by-products is increasing because of limitations in food sources for livestock which result in economic and environmental concerns. Rice straw, a major agricultural by-product, is routinely utilized as a food source for ruminants in many regions of East and South-East Asia (Zhang et al., 2017). In Malaysia, rice straw is one of the most abundant agricultural by-products (Ghazali et al., 2013). However, rice straw has very low nutritive values with low crude protein content and metabolic energy for ruminants. Technologies to create high-quality animal feed from agricultural residues need to be developed. Ensiling is a practical way to utilize water-soluble carbohydrates by lactic acid bacteria (LAB) under anaerobic conditions to produce organic acids such as lactic acid to reduce pH and to inhibit the growth of harmful bacteria resulting in good quality silage (He et al., 2018). Silage feeding is also a way of enhancing livestock production in the tropics especially during periods of inadequate supply of fresh forage. According to the literature, LAB (homofermentative and heterofermentative) which are widely used as inoculants, increased the concentration of lactic acid while lowered the pH and the concentration of NH_3_-N in silage (Silva et al., 2016). Several studies have shown the effectiveness of LAB on the feed quality of rice straw (Zhang et al., 2010; Cao et al., 2013; LIU et al., 2015; Oladosu et al., 2016). Besides, those studies mentioned that adding LAB increased the lactic acid content of silage, increased dry matter digestibility, improved *in vitro* ruminal fermentation parameters and decreased ruminal methane production. However, not all *in vitro* studies have reported reductions in methane production (Contreras-Govea et al., 2011). It has been hypothesized that LAB silage inoculants could reduce methane emissions from ruminants by several modes of action; changes in the chemical composition of the silage, interaction of LAB with rumen microbes and alteration of rumen fermentation (EllIS et al., 2016). Methane, as produced from anaerobic fermentation in the rumen, accounts for 2-12% loss of dietary gross energy in ruminants and is a potent greenhouse gas with a global warming potential 23 times higher than that of carbon dioxide in trapping the heat (Jafari et al., 2018). Therefore, reducing ruminal methane production not only improves the efficiency of nutrient utilization in ruminants but also helps to protect the environment from the negative consequences of global warming.

From the microbiological perspective, some studies indicated that the inclusion of silage alone (Nguyen et al., 2017) as well as silage + LAB inoculant (He et al., 2018) could improve microbial population in the rumen. However, to the best of our knowledge, there is still limited information on the effect of different types of LAB inoculated rice straw silage on microbial population responses. Therefore, the purpose of this experiment was to test the rumen microbial populations and fermentation characteristics as well as testing methane mitigation potential of rice straw silage inoculated with different types of LAB in an *in vitro* condition.

## Materials and Methods

The protocol for the experimental procedures were reviewed and approved by the Animal Care and Use Committee of the University of Putra in Malaysia.

### Isolation, identification and characterization of LAB

Cecal contents from healthy adult, commercial broiler chickens and rumen samples from fistulated male cattle (body weight: 209 kg) were used for the isolation of LAB. 1 gram of each samples were dissolved in 9 ml of peptone water (0.01%) and shaken at 200 rpm for 10 min. Several dilution from each sample (10^−3^ to 10^−7^) were prepared into dilution tube containing peptone water (0.01%). 100 µl of each dilution were transferred into the plate containing MRS Rogosa agar (Oxoid CM 627, Hampshire, UK) as selective medium for LAB (Ebrahimi, 2012). Plates were anarobically incubated at 37 °C for 48h. Several clones were selected from each plate and subcultured for three times. Total of 80 isolates were selected and tested for Gram stain, hydrogen peroxidase and lactic acid production. The LAB strains that actively produced lactic acid were chosen for the molecular identification.

### Molecular identification

DNA of selected LAB was extracted using DNA extraction kit (QIAamp Blood and Tissue Kit, Qiagen, Hilden, Germany). The amplification of 16SrRNA genes were conducted using 27F 5’-AGAGTTTGATCCTGGCTCAG-3’ and 1492R-5’-GGCTACCTTGTTACGACTT-3’primers. The PCR amplification was performed with i-StarTaq DNA polymerase kit (iNtRON Biotechnology, Sungnam, Kyungki-Do, Korea) using 1 μl of a template (10 ng μl^−1^) in 20 μl of reaction solution. Amplification was performed using a BIORAD MyCycler™ thermal cycler with the following program: 1 cycle at 94°C for 4 min, 30 cycles of 94°C for 1 min, 55°C for 30s, 72°C for 2 min and a final extension at 72°C for 5 min. The PCR products were mixed with loading dye and loaded on to a 1.0% SeaKem® GTG® agarose (FMC BioProducts, Rockland ME, USA) containing ethidium bromide, and electrophoresis was carried out at 90 V for 1 h. The PCR products were visualized under UV illumination and excised from the gel and the PCR product was extracted using MEGAquick-spin PCR & Agarose Gel Extraction kit (iNtRON Biotechnology). PCR product was sequenced using forward and reverse primers (1^st^ Base Co., Malaysia). The contig was done for the forward and reverse sequences of each isolates by contig assembly program of Bioedit software and then sequences were analyzed by the Bellerophon and Mallard program to remove chimeric rDNA clones. Approximately 1400 bp segment of the 16S rRNA gene of the isolates were blast using National Center for Biotechnology Information (NCBI) library with the following address: http://blast.ncbi.nlm.nih.gov/Blast.cgi.

### Rice straw ensiling and inoculating procedures

Fresh rice straws used in this experiment were harvested in the fields of the Malaysian Agricultural Research and Development Institute (MARDI) located in Serdang, Selangor, Malaysia (3°00′18.88″N, 101°42′15.05″E). Then, they were chopped to 8 - 10 cm long pieces with a laboratory chopper. Five isolates of LAB (*L. plantarum, L. salivarius, L. reuteri, L. brevis, and S. bovis*) were used for inoculation and the inoculation rate was based on the numbers of colony-forming units per gram in the inoculant powders. The dry matter of chopped rice straw was determined and the inoculants were applied by suspending the appropriate weight of inoculant powder in required amount of water to increase the moisture content of rice straw to 70% and spraying it over 2 kg batches of rice straw and mixed thoroughly. Each treatment contained 10^6^ cfu/g DM of LAB inoculants. The treated rice straw was ensiled in 500mL Scott bottle. There were 3 bottles per inoculant treatment of each of the silages. The silages were stored for 15 and 30 days at room temperature (28 to 32°C). Control silages were also prepared at the same time with sterile water.

### Chemical analyses and fermentation quality for rice straw silage

After 15 and 30 days of ensiling, bottles of the untreated and inoculated silages were opened for analyzing chemical analyses and fermentation quality. 20 g of representative silage were mixed with 180 g sterile water in a laboratory blender (Waring, New Hartford, Conn, USA) for 2 minutes. The extract was filtered through four layers of gauze and no. 1 filter paper (Whatman, Inc., Clifton, NJ). The filtrate extract was used for measuring Dry matter (DM), Crude protein (CP), neutral detergent fiber (NDF), acid detergent fiber (ADF), NH_3_-N, pH, LAB population, lactic acid, and volatile fatty acids (VFA). The DM and CP (total nitrogen × 6.25) were contents determined using method number 934.1 and 990.03 (AOAC, 1990), respectively. NDF and ADF were determined according to Van Soest and coworkers (Van Soest et al., 1991). The concentration of NH_3_-N was determined as described in our previous work (Jafari et al., 2016). The pH was determined using a pH electrode (Mettler-Toledo Ltd., England). Lactic acid and volatile fatty acids were determined using gas-liquid chromatography with Quadrex 007 Series (Quadrex Corporation, New Haven, CT 06525 USA) bonded phase fused silica capillary column (15m, 0.32mm ID, 0.25 µm film thickness) in an Agilent 7890A gas-liquid chromatography (Agilent Technologies, Palo Alto, CA, USA) equipped with a flame ionization detector (FID). The total number of LAB in the silage was determined on MRS Rogosa agar as described above with the plate count method (Ebrahimi, 2012). Colonies were counted from the plates at appropriate dilutions and the number of colony forming units (CFU) was expressed as log10 per gram of rice straw.

### *In vitro* rumen fermentation and digestibility

Two rumen fistulated mature cows were fed (Table 1) at maintenance level on mixed forage and concentrate at 60:40 ratio twice daily. Rumen liquor was collected before the morning feed from both fistulated cows and strained through four layers of muslin gauze into a pre-warmed bottle at 39°C. Treated and untreated rice straw used as substrates. A total of six syringes for each treatment were used for *in vitro* study. The contents of three syringes were used for *in vitro* dry matter digestibility (IVDMD), fermentation parameters and the remaining three syringes were used for rumen microbial population quantification. 500 mg of substrate were weighed into 100 ml calibrated glass syringes. The incubation medium was prepared as described by our previous work (Jafari et al., 2017) and 40 ml was dispensed anaerobically into each syringe. Syringes were incubated at 39 °C for 24 h. *In vitro* gas production was measured in triplicate at 2, 4, 8, 12 and 24. In each incubation run, three blanks were used as blank to correct the values for gas released from the substrates. Cumulative gas production data were fitted in NEWAY Excel Version 5.0 package (Ørskov and McDonald, 1979). The above procedures were conducted in three individual runs. After 24 h of fermentation, IVDMD of substrates was determined by the contents of syringes. The fermentation end products (e.g. pH, NH_3_-N and VFA) and the number of LAB were also determined as described earlier.

**Table 1.** Ingredients and chemical composition of the diets fed to the cows for *in vitro* study

| **Ingredients (g/kg DM)** |  |
| --- | --- |
| Alfalfa Hay | 314.10 |
| Corn, grain | 170.00 |
| Soybean meal | 133.00 |
| Palm kernel cake | 251.10 |
| Rice Bran | 81.80 |
| Sunflower oil | 20.00 |
| Mineral Premix | 5.00 |
| Vitamin Premix | 5.00 |
| Ammonium chloride | 10.00 |
| Limestone | 10.00 |
| **Chemical composition (g/kg DM)** |  |
| DM (%) | 850.20 |
| CP | 208.30 |
| EE | 52.50 |
| NDF | 419.00 |
| ADF | 253.00 |

DM, dry matter; CP, crude protein, EE, ether extract; NDF, neutral detergent fiber; ADF, acid detergent fiber.

### Quantification of rumen microbial population by real-time PCR

The targeted microbes were cellulolytic bacteria such as *Fibrobacter succinogenes, Ruminococcus albus, Ruminococcus flavefaciens*, general bacteria, general anaerobic fungi, total protozoa, total methanogens and total archaea. DNA was extracted from 300 µl of fermented rumen content (fluid and digesta from three syringes) by QIAGEN DNA Mini Stool Kit (QIAGEN, Valencia, CA) according to manufacturer’s recommendations. Then the PCR product was purified using a QIA quick PCR purification kit (QIAGEN, Inc., Valencia, CA) and cloned to the plasmid. The target DNA was quantified by using serial 10-fold dilutions from 10^1^ to 10^8^ DNA copies of the previously quantified DNA purified plasmid. Microorganisms and sequences of the primers used in this study are shown in Table 2.

**Table 2.**
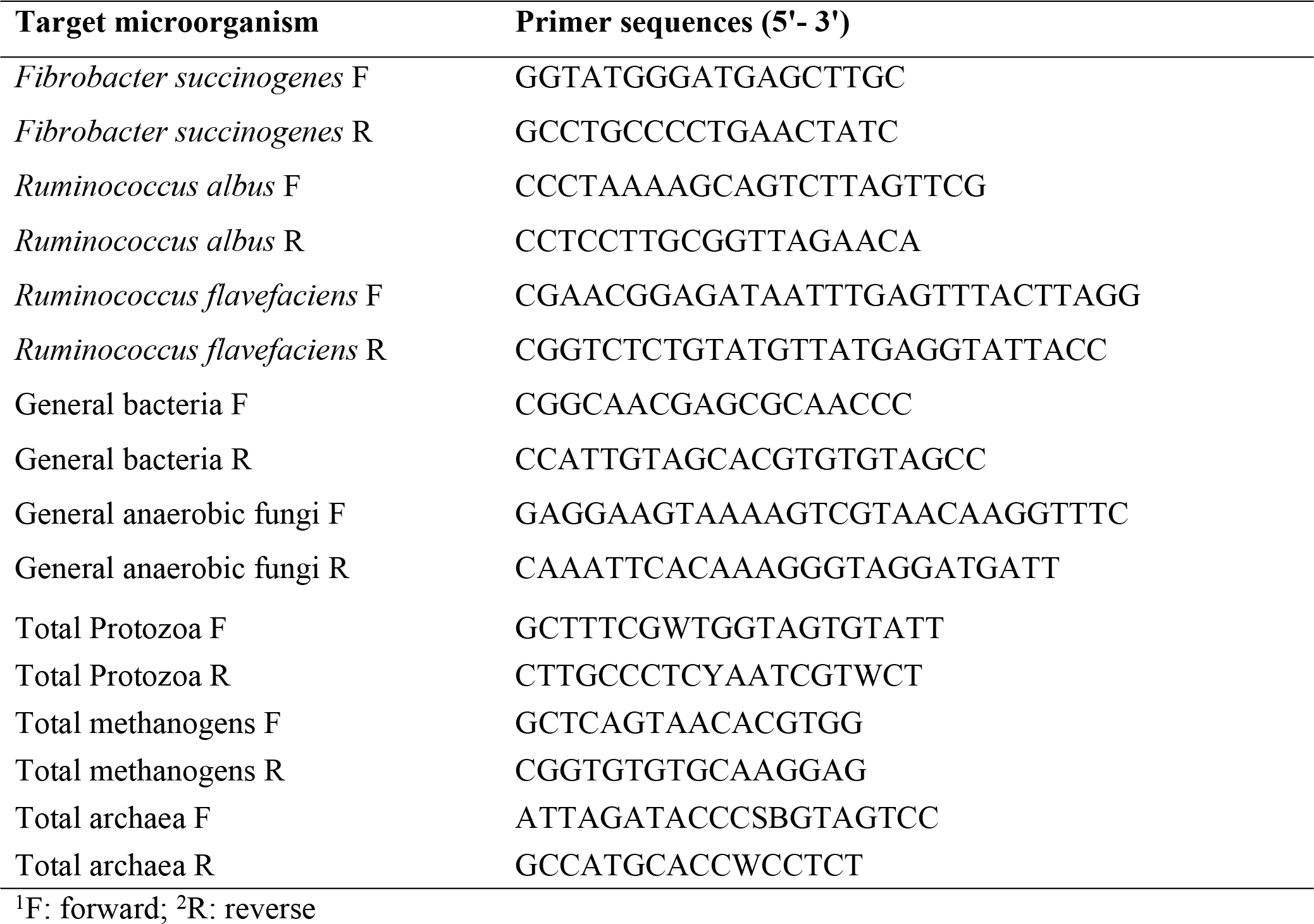
Microorganisms and sequences of the primers used in this study (Ebrahimi 2012).

### Statistical analyses

Data were analyzed using the general linear (GLM) models procedure of SAS (SAS, 2003) in a completely randomized design and the means were compared with Duncan’s Multiple Range test. Differences of P < 0.05 were considered to be significant.

## Results

### Chemical analyses and fermentation quality of rice straw silage

The contents of DM, CP, ether extract, NDF, ADF were affected (P˂0.05) by the treatments (Table 3). The DM contents were numerically decreased as the duration of ensiling increased. The control group had higher DM content as compared with the LAB treatments at 15 and 30 d of ensilage. The CP content was greater in LAB treatments as compared with control (10.9-12.7 *vs* 9.5, respectively). The NDF and ADF of the LAB treatments were less than those of the control (Table 3). However, the gross energy was not affected (P˃0.05) by the treatments. Analysis of sugar in fermented rice straw showed significant decrease (P˂0.05) in the concentration of glucose and fructose among LAB treatments as compared with the control.

**Table 3.** Effect of inoculation of LAB on chemical composition of ensiled rice straw (DM basis)

| Treatments | Day | DM | CP | NDF | ADF | GE | Glucose<sup>1</sup> | Fructose<sup>1</sup> | Xylose<sup>1</sup> |
| --- | --- | --- | --- | --- | --- | --- | --- | --- | --- |
| Control | 15 | 34.8<sup>a</sup> | 9.3<sup>c</sup> | 69.2<sup>a</sup> | 56.2<sup>a</sup> | 15.7<sup>a</sup> | 53.6<sup>a</sup> | 0.4<sup>a</sup> | 0.4 |
|  | 30 | 33.6<sup>a</sup> | 9.8<sup>c</sup> | 70.2<sup>a</sup> | 56.8<sup>a</sup> | 15.9<sup>a</sup> | 48.2<sup>b</sup> | 0.4<sup>a</sup> | 0.4 |
| *L. plantarum* | 15 | 30.5<sup>bc</sup> | 11.8<sup>b</sup> | 66.8<sup>c</sup> | 53.3<sup>b</sup> | 15.5<sup>a</sup> | 35.9<sup>c</sup> | 0.3<sup>ab</sup> | 0.4 |
|  | 30 | 29.2<sup>c</sup> | 12.7<sup>a</sup> | 64.1<sup>d</sup> | 49.4<sup>c</sup> | 14.8<sup>ab</sup> | 12.2<sup>h</sup> | 0.1<sup>c</sup> | 0.3 |
| *L. salivarius* | 15 | 33.2<sup>a</sup> | 11.2<sup>b</sup> | 68.9<sup>a</sup> | 53.8<sup>b</sup> | 15.7<sup>a</sup> | 36.1<sup>c</sup> | 0.3<sup>ab</sup> | 0.4 |
|  | 30 | 32.8<sup>ab</sup> | 12.1<sup>ab</sup> | 66.6<sup>c</sup> | 51.9<sup>c</sup> | 15.5<sup>a</sup> | 25.4<sup>f</sup> | 0.2<sup>b</sup> | 0.4 |
| *L. reuteri* | 15 | 32.5<sup>ab</sup> | 10.9<sup>bc</sup> | 67.6<sup>b</sup> | 53.7<sup>b</sup> | 15.9<sup>a</sup> | 32.7<sup>d</sup> | 0.3<sup>ab</sup> | 0.4 |
|  | 30 | 31.3<sup>b</sup> | 10.9<sup>bc</sup> | 66.6<sup>c</sup> | 52.5<sup>bc</sup> | 15.0<sup>a</sup> | 24.6<sup>f</sup> | 0.2<sup>b</sup> | 0.3 |
| *L. brevis* | 15 | 33.4<sup>a</sup> | 10.6<sup>bc</sup> | 66.9<sup>c</sup> | 52.9<sup>bc</sup> | 15.3<sup>a</sup> | 33.2<sup>d</sup> | 0.3<sup>ab</sup> | 0.4 |
|  | 30 | 32.4<sup>ab</sup> | 10.9<sup>bc</sup> | 66.6<sup>c</sup> | 50.6<sup>c</sup> | 15.1<sup>a</sup> | 30.4<sup>e</sup> | 0.2<sup>b</sup> | 0.3 |
| *S. bovis* | 15 | 32.6<sup>ab</sup> | 12.4<sup>a</sup> | 67.5<sup>b</sup> | 52.1<sup>bc</sup> | 15.7<sup>a</sup> | 34.6<sup>cd</sup> | 0.3<sup>ab</sup> | 0.3 |
|  | 30 | 31.5<sup>b</sup> | 12.5<sup>a</sup> | 64.8<sup>d</sup> | 50.8<sup>c</sup> | 15.3<sup>a</sup> | 19.7<sup>g</sup> | 0.2<sup>b</sup> | 0.3 |
| SEM | - | 0.60 | 0.43 | 0.46 | 0.97 | 0.65 | 0.83 | 0.05 | 0.08 |

<sup>1</sup>Unit: mg/g
DM, dry matter; CP, crude protein; NDF, neutral detergent fiber; ADF, acid detergent fiber; GE, gross energy.
Means in each column with different superscripts are significantly different (P<0.05).
SEM, standard error of the mean.

The value of pH decreased in all treatments except for control as the duration of ensiling increased (from 15 d to 30 d). The LAB treatment groups showed the lowest pH value as compared with control throughout the ensiling period with pH values between 4.3-5.3. Lactic acid content (mM) increased from days 15 to 30 among all treatments; however, LAB treatments were significantly higher than that of control. Among the LAB treatments, *L. plantarum* and *S. bovis* had the highest lactic acid content at 30 d of ensilage (36.9 and 35.7, respectively). The acetic acid and propionic acid contents of all treatments increased with the increase in duration of ensiling. Again, *L. plantarum* and *S. bovis* showed the greatest values for acetic and propionic acids at 30 d of ensilage (24.1 and 2.9 *vs* 22.5 and 2.5 mM, respectively). Butyric acid content showed a decreasing trend among the treatments as the duration of ensiling increased, with the highest value for the control (5.5 and 4.6 mM at 15 and 30 d of ensilage, respectively). As compared with the control, LAB treatments didn’t show significant differences (P˃0.05) in terms of NH_3_-N concentration (average: 0.049%). The analysis of the LAB content (log cfu/g) showed that the LAB treatments exhibited a significant (P < 0.05) difference and increase as compared with control (Table 4).

**Table 4.** Effects of inoculation of LAB on fermentation quality of ensiled rice straw

|  | D<br>a<br>y | pH | Lactic<br>acid<br>(mM) | Acetic<br>acid<br>(mM) | Propioni<br>c acid<br>(mM) | Butyric<br>acid<br>(mM) | NH<sub>3</sub>-N<br>(%) | LAB<br>(log<br>cfu/g) |
| --- | --- | --- | --- | --- | --- | --- | --- | --- |
| Control | 1<br>5<br>3<br>0 | 5.6<sup>a</sup><br>5.6<sup>a</sup> | 5.1<sup>f</sup><br>9.4<sup>e</sup> | 10.5<sup>e</sup><br>11.6<sup>e</sup> | 1.3<sup>b</sup><br>1.6<sup>b</sup> | 5.5<sup>a</sup><br>4.6<sup>a</sup> | 0.05<br>0.05 | 5.2<sup>c</sup><br>5.8<sup>c</sup> |
| *L. plantarum* | 1<br>5<br>3<br>0 | 5.2<sup>ab</sup><br>4.4<sup>b</sup> | 19.6<sup>c</sup><br>36.9<sup>a</sup> | 20.8<sup>b</sup><br>24.1<sup>a</sup> | 1.9<sup>ab</sup><br>2.9<sup>a</sup> | 1.7<sup>cd</sup><br>1.7<sup>cd</sup> | 0.04<br>0.05 | 6.6<sup>bc</sup><br>8.8<sup>a</sup> |
| *L. salivarius* | 1<br>5<br>3<br>0 | 5.4<sup>a</sup><br>4.8<sup>b</sup> | 14.6<sup>d</sup><br>22.4<sup>bc</sup> | 13.4<sup>d</sup><br>19.1<sup>bc</sup> | 1.2<sup>b</sup><br>1.3<sup>b</sup> | 3.3<sup>b</sup><br>2.4<sup>b</sup> | 0.05<br>0.05 | 6.5<sup>bc</sup><br>7.3<sup>b</sup> |
| *L. reuteri* | 1<br>5<br>3<br>0 | 5.5<sup>a</sup><br>4.8<sup>b</sup> | 15.5<sup>d</sup><br>26.6<sup>b</sup> | 14.2<sup>d</sup><br>17.4<sup>c</sup> | 1.5<sup>b</sup><br>2.1<sup>ab</sup> | 2.2<sup>c</sup><br>2.1<sup>c</sup> | 0.05<br>0.06 | 6.4<sup>bc</sup><br>7.1<sup>b</sup> |
| *L. brevis* | 1<br>5<br>3<br>0 | 5.3<sup>a</sup><br>4.9<sup>b</sup> | 16.9<sup>c</sup><br>24.1<sup>b</sup> | 14.4<sup>d</sup><br>18.1<sup>c</sup> | 1.4<sup>b</sup><br>1.7<sup>b</sup> | 2.6<sup>b</sup><br>2.1<sup>c</sup> | 0.05<br>0.05 | 6.6<sup>bc</sup><br>7.3<sup>b</sup> |
| *S. bovis* | 1<br>5<br>3<br>0 | 5.3<sup>a</sup><br>4.3<sup>b</sup> | 19.9<sup>c</sup><br>35.7<sup>a</sup> | 19.5<sup>bc</sup><br>22.5<sup>ab</sup> | 1.6<sup>b</sup><br>2.5<sup>a</sup> | 1.7<sup>cd</sup><br>1.2<sup>d</sup> | 0.04<br>0.05 | 6.8<sup>bc</sup><br>8.2<sup>ab</sup> |
| SEM | - | 0.31 | 1.14 | 0.88 | 0.53 | 1.01 | 0.004 | 0.48 |

Means in each column with different superscripts are significantly different (P<0.05).
SEM, standard error of the mean.

### *In vitro* rumen fermentation characteristics, methane production and DM digestibility

According to the data of *in vitro* (Table 5), LAB treatments had less (P˂0.05) gas production at 24 h of fermentation as compared with control. Conversely, coefficient of degradable *B* fraction was greater in LAB treatments especially for *L. plantarum* and *S. bovis* (at 30 d of ensilage) as compared with control. However, coefficients of rapidly degradable *a* fraction and *c* (degradation rate of degradable b fraction) were not affected (P˃0.05) by the treatments. The LAB treatments especially for *L. plantarum* and *S. bovis* at 30 d of ensiling had greater (P˂0.05) amounts of IVDMD as compared with control. Total VFA and acetic acid was also greater (P˂0.05) among LAB treatments. The concentration of NH_3_-N and pH were almost similar among the treatments with no significant difference (P˃0.05). Methane production and methane/total gas significantly (p<0.05) decreased between LAB treatments compared with control groups. *L. plantarum* at 15 and 30 d of ensiling exhibited respectively 46% and 48% of CH_4_ reduction as compared with control.

**Table 5.** Effects of inoculation of LAB on *in vitro* rumen fermentation

| Treatments | Day | DMD | Total gas | pH | NH<sub>3</sub>-N | Total VFA (mM) | Acetic acid | Propionic acid | CH<sub>4</sub> | CH<sub>4</sub>/total gas | a | b | c | a+b |
| --- | --- | --- | --- | --- | --- | --- | --- | --- | --- | --- | --- | --- | --- | --- |
| Control | 15 | 21.4<sup>f</sup> | 45.5<sup>a</sup> | 6.9 | 14.3 | 65.6<sup>g</sup> | 45.5<sup>g</sup> | 13.6<sup>a</sup> | 7.8<sup>a</sup> | 0.17<sup>a</sup> | 7.0 | 43.0<sup>cd</sup> | 0.05 | 50.0<sup>d</sup> |
|  | 30 | 22.2<sup>f</sup> | 45.0<sup>a</sup> | 6.9 | 15.6 | 67.5<sup>ef</sup> | 44.5<sup>g</sup> | 12.7<sup>ab</sup> | 7.9<sup>a</sup> | 0.18<sup>a</sup> | 6.0 | 44.0<sup>c</sup> | 0.05 | 50.0<sup>d</sup> |
| *L. plantarum* | 15 | 25.4<sup>de</sup> | 42.0<sup>bc</sup> | 6.9 | 15.3 | 74.7<sup>c</sup> | 46.1<sup>fg</sup> | 12.4<sup>b</sup> | 4.2<sup>c</sup> | 0.10<sup>cd</sup> | 6.5 | 44.0<sup>c</sup> | 0.05 | 50.5<sup>cd</sup> |
|  | 30 | 29.4<sup>a</sup> | 37.5<sup>c</sup> | 6.9 | 15.9 | 79.7<sup>a</sup> | 54.4<sup>a</sup> | 12.9<sup>ab</sup> | 4.1<sup>c</sup> | 0.11<sup>cd</sup> | 7.7 | 48.5<sup>a</sup> | 0.05 | 56.2<sup>a</sup> |
| *L. salivarius* | 15 | 22.4<sup>f</sup> | 43.5<sup>b</sup> | 6.8 | 15.2 | 68.8<sup>e</sup> | 47.0<sup>f</sup> | 12.4<sup>b</sup> | 6.1<sup>b</sup> | 0.14<sup>b</sup> | 6.5 | 43.5<sup>c</sup> | 0.05 | 50.0<sup>d</sup> |
|  | 30 | 26.4<sup>cd</sup> | 41.0<sup>c</sup> | 6.8 | 15.8 | 76.1<sup>b</sup> | 49.7<sup>d</sup> | 13.0<sup>ab</sup> | 5.9<sup>b</sup> | 0.14<sup>b</sup> | 6.5 | 45.5<sup>b</sup> | 0.05 | 52.0<sup>c</sup> |
| *L. reuteri* | 15 | 22.4<sup>f</sup> | 40.5<sup>c</sup> | 6.8 | 14.9 | 70.6<sup>d</sup> | 46.7<sup>f</sup> | 13.3<sup>a</sup> | 4.5<sup>c</sup> | 0.11<sup>cd</sup> | 6.5 | 43.5<sup>c</sup> | 0.05 | 50.0<sup>d</sup> |
|  | 30 | 26.4<sup>cd</sup> | 40.0<sup>cd</sup> | 6.8 | 15.5 | 74.5<sup>c</sup> | 51.6<sup>c</sup> | 13.2<sup>a</sup> | 4.6<sup>c</sup> | 0.12<sup>c</sup> | 6.0 | 45.7<sup>b</sup> | 0.05 | 51.7<sup>c</sup> |
| *L. brevis* | 15 | 22.1<sup>f</sup> | 43.0<sup>b</sup> | 6.8 | 15.3 | 67.2<sup>ef</sup> | 48.3<sup>de</sup> | 12.5<sup>b</sup> | 5.2<sup>b</sup> | 0.12<sup>c</sup> | 6.25 | 42.5<sup>cd</sup> | 0.05 | 48.7<sup>e</sup> |
|  | 30 | 27.4<sup>bc</sup> | 42.0<sup>bc</sup> | 6.8 | 15.6 | 78.3<sup>ab</sup> | 52.7<sup>b</sup> | 13.6<sup>a</sup> | 5.4<sup>b</sup> | 0.13<sup>bc</sup> | 7.5 | 46.0<sup>b</sup> | 0.05 | 53.5<sup>bc</sup> |
| *S. bovis* | 15 | 24.4<sup>ef</sup> | 40.5<sup>c</sup> | 6.8 | 15.1 | 74.5<sup>c</sup> | 49.2<sup>d</sup> | 13.8<sup>a</sup> | 5.7<sup>b</sup> | 0.14<sup>b</sup> | 7.0 | 44.5<sup>bc</sup> | 0.05 | 51.5<sup>c</sup> |
|  | 30 | 28.4<sup>ab</sup> | 39.5<sup>cd</sup> | 6.8 | 15.7 | 78.9<sup>a</sup> | 53.5<sup>ab</sup> | 13.3<sup>a</sup> | 5.2<sup>b</sup> | 0.13<sup>bc</sup> | 6.75 | 47.5<sup>a</sup> | 0.05 | 54.2<sup>b</sup> |
| SEM | - | 0.68 | 0.84 | 0.03 | 0.48 | 1.07 | 0.48 | 0.32 | 0.26 | 0.004 | 0.17 | 1.05 | 0.01 | 0.92 |

a, b, c, and a+b were calculated from exponential equation $p=a+b(1-e^{-ct})$ .
a = gas production from the immediately soluble fraction, b = gas production from the insoluble fraction, c = gas production rate constant for the insoluble fraction (b),
(a + b) = potential extent of gas production.
DMD, dry matter digestibility; VFA, volatile fatty acid.
Means in each column with different superscripts are significantly different ( $P<0.05$ ).

### *In vitro* rumen microbial populations

LAB treatments had greater (P˂0.05) total bacteria and fungi at 24 h of fermentation as compared with control (Table 6). Conversely, control had greater (P˂0.05) total protozoa, methanogens and archaea at the end of *in vitro* fermentation. *Butyrivibrio fibrisolvens* and *Ruminococcus flavafaciences* was lower for control as compared with LAB treatments at 15 and 30 days of silage. Especially, *L. plantarum* and *S. bovis* at 30 d of silage had the greatest populations of mentioned bacteria. *Fibrobacter succinogenes* was also almost similar among LAB treatments but higher (P˂0.05) than control group.

**Table 6.**
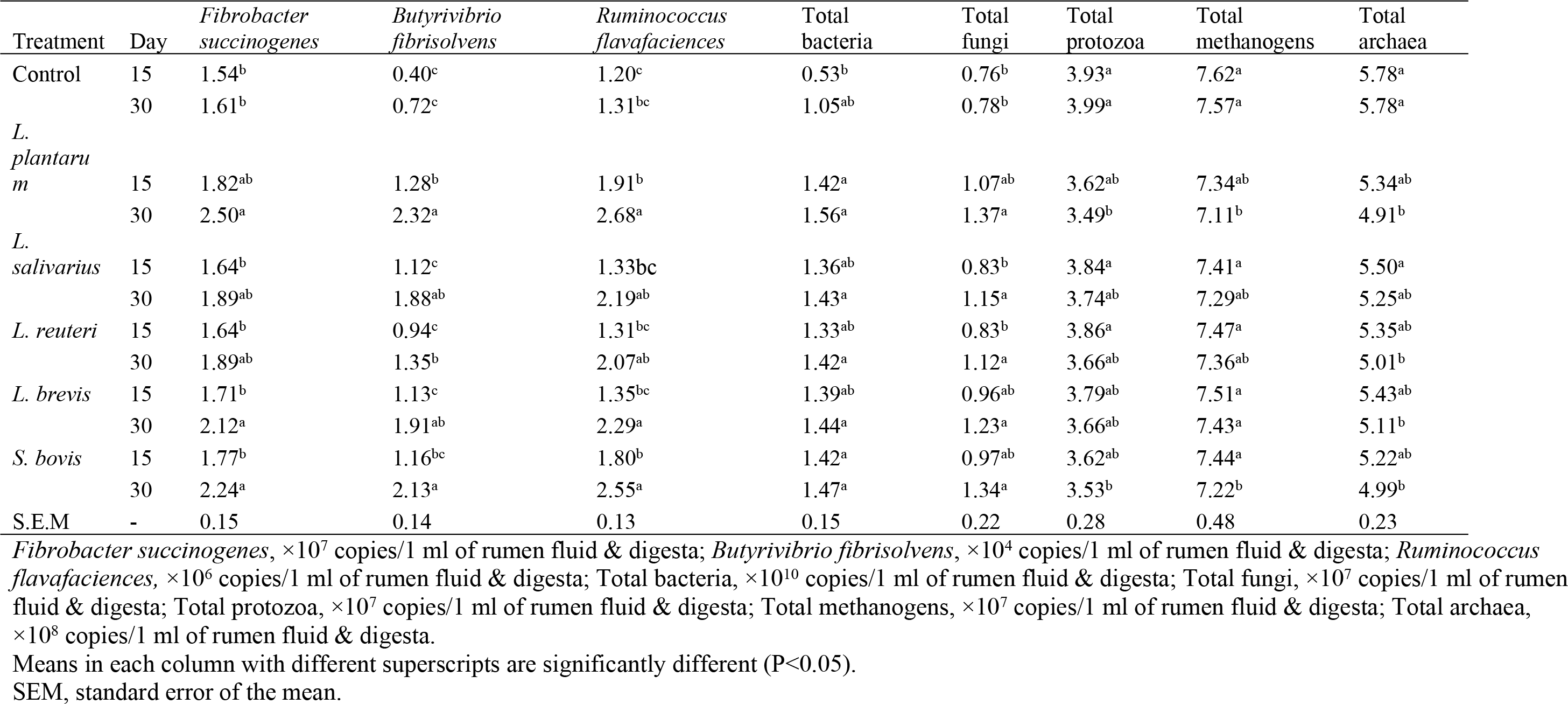
Effects of inoculation of LAB on *in vitro* rumen microbial populations

## Discussion

### Chemical composition and fermentation characteristics

Inoculating the different types of LAB for ensiled rice straw increased CP content in the current study. Our results were consistent with the results of Liu et al. (2015) which evaluated the effects of fermentation using LAB culture broth on the feed quality of rice straw. However, our results are contradictory to their results in terms of NH_3_-N, NDF and ADF contents. The LAB treatments in the current study did not affect NH_3_-N concentration after 15 and 30 days of ensiling. High concentration of NH_3_-N is the results of excessive breakdown of protein during fermentation which lowers silage quality (Jafari et al., 2018). Lower NDF and ADF contents among the LAB treatments compared with the control group in this study could be the result of the lower level of heat damage on protein which improves energy content (Saha et al., 2010) as shown in Table 3. Chen et al. (2019) mentioned that lower NDF content in silages could also be due to the loss of hemicellulose occurred in the ensiling process. This loss could be due to a combination of enzymatic and acid hydrolysis of the more digestible cell wall fractions during the fermentation. DM is the remaining materials after the removal of water and contains the main nutrients for animal consumption. Ensilage of the forage will mostly result in the DM loss which occurs during the fermentation. In the current study, inoculations of different LAB decreased the DM loss which could be due to inhibiting the *clostridia* and aerobic bacteria (Ni et al., 2015). The lack of DM loss in our study was also consistent with the application of LAB isolated from forage paddy rice silage in China (Ni et al., 2015).

The previous studies showed that bacterial inoculation of silage could convert the composition of cell-wall carbohydrates into organic acids and cause a decrease in pH during fermentation (Baek et al., 2017). In the current study, pH decreased with increase in the duration of ensiling among the LAB treatments (especially for *L. plantarum* and *S. bovis*) which was consistent with those in the literature. Silage pH (the lower the better) is one of the main factors depicting the extent of fermentation and quality of ensiled forage (Chen et al., 2019). The lower pH among LAB treatments (4.99) *VS* control (5.6) suggests good fermentation. Consistently, Kim et al. (2017) indicated that *L. plantarum* inoculant for fresh rice straw silage decreased the pH, acetic acid, NH_3_-N, and butyric acid contents (Kim et al., 2017). However, in our study, LAB treatments improved acetic acid with no effect on butyric acid content. A high concentration of butyric acid is the sign of protein degradation and DM loss as well as energy wastage (Oladosu et al., 2016). Kim et al. (2017) also concluded that adding *L. plantarum* could improve the fermentation quality and feed value of rice straw silage. Inoculation of the mixture of corn steep liquor and air-dried rice straw with homo fermentative (*L. plantarum*) and hetero-fermentative (*L. plantarum, Lactobacillus casei*, and *Lactobacillus buchneri*) LAB significantly increased the concentration of acetic acid and lactic acid compared with the control in a study conducted in China (Li et al., 2016). Our results were also consistent with that study in terms of increased acetic acid and lactic acid contents. High concentration of lactic acid results in lower pH (as observed in this study) which inhibits the growth and activities of undesirable bacteria during silage (Oladosu et al., 2016). Acetic acid also possesses antifungal activity which reduces the spoilage of organisms in ensiled mass and improves the fermentation quality of silages. Zhang et al. (2010) mentioned that chopping rice straw before ensiling could enhance the lactic acid concentration and total VFA content. The improved criteria observed in our study could also be due to chopping the rice straw before ensiling. Li et al. (2016) also demonstrated that homo fermentative and hetero-fermentative LAB could effectively improve the fermentation quality of the silage. Rice straw, a by-product of rice production which could be abundantly found in Southeast Asia which is the most important rice-producing region in the world (Zhang et al., 2010). Thus, by improving the nutritive value of this by-product through processes such as ensiling and inoculating beneficial microbes, farmers could overcome the limitations of feed sources in many parts of the tropics.

### *In vitro* rumen fermentation characteristics, methane production and DM digestibility

Some studies have reported the effectiveness of LAB inoculation on *in vitro* ruminal fermentation characteristics (Zhang et al., 2016; Baek et al., 2017; Zhang et al., 2017). Lack of effect on rumen pH and NH_3_-N after 15 and 30 days of ensiling among the LAB treatments in this study was contrary to the results of Zhang et al. (2010). They reported that three levels of LAB inoculants (LAB; 2×10^5^, 3×10^5^ and 4×10^5^ cfu/g fresh forage) on rice straw (whole and chopped rice straw) silage decreased pH, NH_3_-N and acetic acid concentrations in Holstein dairy cows. Our results were consistent with theirs in terms of total VFA and propionic acid concentrations which showed respectively increase and decrease among the LAB treatments. Zhang et al. (2010) also concluded that the chopping process and LAB addition improved the silage quality of rice straw, and its partial substitution with corn silage could lower the cost of the dairy cow ration with no negative effects on lactation performance. Supplementing rice straw and sugar beet leaf silage treated with lactic acid bacteria enhanced performance and productivity of lactating Frisian cows in an *in vivo* study (El Tawab et al., 2017). Another *in vivo* study showed improved fermentation quality, as well as improved digestibility of feed components after feeding wethers with urea treated rice straw silage with LAB (Fang et al., 2012). In this study, the LAB treatments especially for *L. plantarum* and *S.bovis* showed the highest IVDMD and the lowest methane production. Different results obtained among variant types of LAB in this study were consistent with Ellis et al. (2016) which showed that organic matter digestibility, gas and methane production vary with type of LAB added and type of substrate incubated. Our results were consistent with the previous studies and the results of Cao et al. (2013) in which vegetable residue silage inoculated with *L. plantarum* showed the highest IVDMD and lowest methane production. Methane is a by-product of the anaerobic fermentation of dietary carbohydrates in the rumen, and methanogenesis possesses a biological regulatory mechanism for animal health (Chen et al., 2017). However, Jafari et al. (2016) mentioned that methane formation is a contributing factor for the atmospheric burden of green-house gases, which is linked to the global warming and climate change as well as a significant energy loss for animal due to the exit of carbon.

### *In vitro* rumen microbial populations

The growing public concern over the widespread use of antibiotics in livestock production and the emergence of antibiotic-resistant bacteria has stimulated interest in developing alternatives that promote animal performance and health. One potential alternative is the use of direct-fed microbials as feed additives to thrive in the gastrointestinal tract and prevent the establishment of pathogens (Jiao et al., 2017). LAB as a particular type of direct-fed microbials as well as LAB silage inoculants has exerted probiotic effects resulted in improvement in ruminant performance (Weinberg et al., 2016). In the current study, microbial populations were affected by the LAB treatments. *Fibrobacter succinogenes*, *Ruminococcus flavefaciens* and *Ruminococcus albus* which are the most predominant cellulolytic bacterial species in ruminants were highest among the LAB treatments. Jiao et al. (2016) reported about the beneficial effect of LAB on fiber digestion which could be due to the competition with the efficient lactate-producing rumen microorganisms on essential compounds. This competition might reduce the rate of lactate production by rumen bacteria which results in higher activity of cellulolytic rumen populations. Our results were in agreement with the previous study (Nguyen et al., 2017). According to Nguyen et al. (2017), dairy steers receiving rice straw and *Leucaena* silage enhanced rumen microbial population (especially cellulolytic), and fungal population as well. They mentioned a decrease in the protozoal populations by the increase in the level of *Leucaena* silage. We also found decreases in the protozoal and methanogen populations by the LAB treatments. Jafari et al. (2018) reported that protozoa can provide electrons as a source of H_2_ to the methanogens, and hence, antiprotozoal effects of feedstuff could decrease methane production by methanogens attached to protozoa. Moreover, fungal populations were increased in our studies among the LAB treatment groups. Nguyen et al. (2017) indicated that there was an increase in the numbers of fungal when protozoa have been removed from the rumen. They also mentioned that *Leucaena* silage could provide adequate nitrogen source for microbial growth leading to the increase in the bacterial population which could be the case for our result’s. Consistent to our study, total mixed rations containing corn silage and/or grass silage increased total bacteria and *Fibrobacter succinogenes* in dairy cows (Lengowski et al., 2016). *B. fibrisolvens* which are involved in rumen fatty acid biohydrogenation were greater among LAB treatments in this study. Conjugated linoleic acid which has beneficial biological effects in animal models is formed as an intermediate during biohydrogenation of linoleic acid to stearic acid in the rumen by mainly *B. fibrisolvens* and other rumen bacteria (Ebrahimi et al., 2018).

### Conclusions

In conclusion, inoculation of Lactobacillus (10^6^ g^−1^ DM) in rice straw silage improved the silage quality (e.g high CP content) and fermentation characteristic (e.g. increase in production of lactic acid and acetic acid) in the silage. Among inoculated LAB, *L. plantarum* and *S. bovis* were found to be more potent for the fermentation. *In vitro* rumen digestibility test showed higher rumen digestibility, higher VFA production and lower methane production in the rice straw fermented with LAB particularly with *L. plantarum* and *S. bovis.* Moreover, analysis of rumen microbial population showed significant increases in the populations of cellulolytic bacterial (*Fibrobacter succinogenes*, *Butyrivibrio fibrisolvens* and *Ruminococcus flavafaciences*), protozoa, methanogens and archaea among the LAB treatments as compared with control. Overall, *L. plantarum, S. bovis* were found to be more promising to be applied in rice straw fermentation; however, *in vivo* experiments need to confirm these results.

## Acknowledgment

The authors would like to thank the faculty of Veterinary Medicine, University of Putra in Malaysia as well as Ratchadapisek Somphot Fund for Postdoctoral Fellowship in Chulalongkorn University.

